# Insight into the resistome and quorum sensing system of a divergent *Acinetobacter pittii* isolate from an untouched site of the Lechuguilla Cave

**DOI:** 10.1101/745182

**Authors:** Han Ming Gan, Peter Wengert, Hazel A. Barton, André O. Hudson, Michael A. Savka

**Affiliations:** Centre for Integrative Ecology, School of Life and Environmental Sciences, Deakin University, Geelong 3220, Victoria, Australia; Deakin Genomics Centre, Deakin University, Geelong 3220, Victoria, Australia; School of Science, Monash University Malaysia, Bandar Sunway, 47500 Petaling Jaya, Selangor, Malaysia; Thomas H. Gosnell School of Life Sciences, Rochester Institute of Technology, Rochester, NY, USA; Department of Biology, University of Akron, Akron, Ohio, USA

**Keywords:** *Acinetobacter*, quorum sensing, antibiotic resistance

## Abstract

*Acinetobacter* are Gram-negative bacteria belonging to the sub-phyla Gammaproteobacteria, commonly associated with soils, animal feeds and water. Some members of the *Acinetobacter* have been implicated in hospital-acquired infections, with broad-spectrum antibiotic resistance. Here we report the whole genome sequence of LC510, an *Acinetobacter* species isolated from deep within a pristine location of the Lechuguilla Cave. Pairwise nucleotide comparison to three type strains within the genus *Acinetobacter* assigned LC510 as an *Acinetobacter pittii* isolate. Scanning of the LC510 genome identified two genes coding for *β*-lactamase resistance, despite the fact that LC510 was isolated from a portion of the cave not previously visited by humans and protected from anthropogenic input. The ability to produce acyl-homoserine lactone (AHL) signal in culture medium, an observation that is consistent with the identification of the *luxI* and *luxR* homologs in its genome, suggests that cell-to-cell communication remains important in an isolated cave ecosystem.

## Introduction

The identification and functionality of antibiotic-resistant and quorum sensing genes from bacteria isolated from pristine environments (areas previously not visited by humans) have raised questions about their origins and natural functions in the environment (1, 2). For example, several bacterial strains isolated from an extremely isolated, hyper-oligotrophic underground ecosystem, Lechuguilla Cave, were shown to harbor antibiotic-resistant genes, including nine previously unrecognized mechanisms of antibiotic resistance (1). In 2014, a total of 93 LC strains (33% Gram-positive and 63% Gram-negative) were reported by Bhullar *et al* and were phylogenetically classified based on sequencing of the 16S rRNA gene (3). In recent years, the taxonomic assignment of some LC strains has been revised mostly at the species level following whole-genome sequencing and genome-based phylogeny (4, 5). In addition, new *luxI* homologs have also been identified in the sequenced strains following genome annotation (4, 5).

Of the 93 LC strains reported, LC510 stood out due to its initial species designation as *Acinetobacter calcoaceticus* that is associated with nosocomial infections. LC510 was isolated from a site deep within the Capitan Formation proximal to the region named “Deep Secrets” at a depth below the surface of approximately 400 m (3). Although most members of *Acinetobacter* are found in soils, waters and occasionally animal feeds, some *Acinetobacter* species are known to infect humans with broad-spectrum antibiotic resistance and such environmental isolates may serve as a reservoir for additional resistance determinants (6, 7). In this study, we characterized LC510 using whole-genome sequencing, biochemical assays, and bioinformatic tools, providing insights into its taxonomic affiliation, resistome and quorum sensing potential.

## Materials and Methods

### DNA extraction and whole-genome sequencing

The isolation, antibiotic characterization and 16S rRNA gene-based identification of strain LC510 have been described previously (3). For gDNA extraction, a single plate colony of LC510 was inoculated into 50 mL of sterile ½ strength tryptic soy broth (TSB) and grown overnight at 30°C with shaking at 150 rpm. The overnight culture was pelleted by centrifugation at 10,000 × g for 10 minutes. Genomic DNA (gDNA) extraction was performed on the pelleted cells using the QIAam DNA Mini kit (Qiagen, Germany) according to the manufacturer’s instructions. The purified gDNA was quantified using the Qubit BR Assay (Invitrogen, Santa Clara, CA, USA) and normalized to 0.2 ng/µL for Nextera XT library preparation (Illumina, San Diego, CA, USA). The constructed library was sequenced on an Illumina MiSeq (2 × 151 bp run configuration) located at the Monash University Malaysia Genomics Facility that routinely sequences metazoan mitogenomes (8-10) and occasionally viral and microbial genomes (11, 12) with no prior history of processing any member from the genus *Acinetobacter*, or more broadly the family Moraxellaceae.

### *De novo* assembly and genome-based species classification

Raw paired-end reads were adapter-trimmed using Trimmomatic v0.36 (13) followed by *de novo* assembly using Unicycler v0.4.7 (default setting with minimum contig length set to 500 bp) (14). We then used Jspecies v1.2.1 (15) to calculate the pairwise average nucleotide identity (ANI) of LC510 against the type strain genomes of *Acinetobacter pittii* (WGS Project: <u>BBST01</u>), *Acinetobacter lactucae* (WGS Project: <u>LRPE01</u>) and *Acinetobacter calcoaceticus* (WGS Project: <u>AIEC01</u>).

### Genome annotation and detection of antibiotic resistance genes

Genome annotation used Prokka v1.13 (16) and the predicted protein-coding genes were used as the input for Abricate v0.8.7 (https://github.com/tseemann/abricate) to search for antibiotic resistance genes against the ResFinder database (minimum query length coverage and nucleotide identity of 90%) (17). The alignment of *bla*_*OXA*_ proteins used MUSCLE followed by maximum likelihood tree construction with FastTree2 (1000 (18).

### Acyl-homoserine lactone bioassay

A single ½ strength tryptic soy agar plate colony of LC510 was inoculated into 50 mL of sterile ½ strength tryptic soy broth and grown overnight at 30°C with shaking at 150 rpm. An equal volume of ethyl acetate (EtOAc) was added to the culture followed by shaking at 50 rpm on an orbital shaker for one hour. The EtOAc layer (upper layer) containing the extracted AHL was evaporated to dryness with a vacuum concentrated and resuspended in fresh EtOAc to make a 20× concentrated extract. Then, 25 µl of the extract was spotted (2 µl/transfer) onto a reverse-phase thin layer chromatography silica gel 60 RP-18 sheet (Merck, Kenilworth, NJ, USA). In addition to the LC510 AHL extract, six synthetic AHL standards were also spoted in separate lanes for comparison. The chromatography was carried-out with a 70%:30% methanol:water mobile phase. The TLC was subsequently dried and overlaid with 1X AB agar medium containing TraR-dependent *lacZ Agrobacterium tumefaciens* reporter strain and X-gal as previously described (19).After an overnight incubation at 30°C, visualization and identification of the AHL separated on the TLC were carried out.

### *In-silico* analysis of the homoserine lactone synthase gene

Proteins were scanned using HMMsearch v3.1b1 with an E-value cutoff of 1E-5 for the presence of Pfam profile PF00765 (https://pfam.xfam.org/family/Autoind_synth) that contains the probabilistic model used for the statistical inference of LuxI-type family of autoinducer synthases (20, 21). The gene organization of contigs containing the *luxI* and *luxR* homolog was visualized using Easyfig (BLASTn setting) (22).

## Results and Discussion

### Genome statistics and taxonomic assignment

The assembled LC510 genome was contained in 115 contigs (N_50_ length of 66.9 kb) with a total length and GC content of 3,767,126 bp and 38.63%, respectively. Based on 16S rRNA gene identification, strain LC510 was previously assigned to the species *Acinetobacter calcoaceticus* (See Table S2 in (3)). However, it only exhibited a pairwise ANI of 90% to *Acinetobacter calcoaceticus* DSM30006^⊤^, a value that is far below the established threshold required for species assignment (15). Expanding the ANI calculation to other closely related species of *A. calcoaceticus* showed that strain LC510 should instead be assigned to the species *A. pittii*, given its >95% pairwise ANI to *A. pittii* DSM25618^⊤^ and *A. pittii* PHEA-2 (Figure 1).

**Figure 1.**
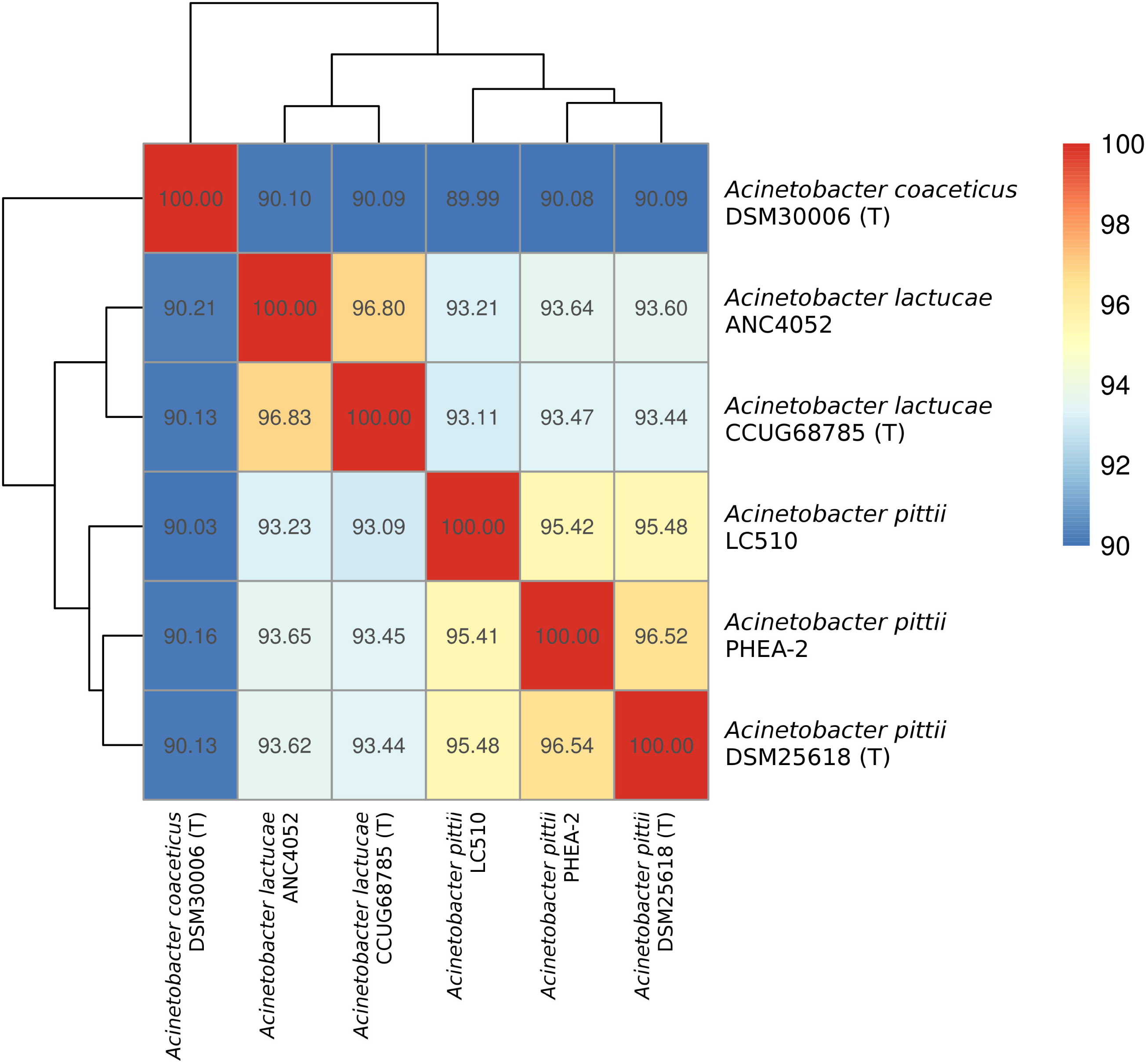
Heatmap showing the clustering of *Acinetobacter* isolates based on their pairwise average nucleotide identity as indicated by the numerical values in the boxes.

### Identification of *beta*-lactamase-producing genes

LC510 was previously shown to exhibit resistance against ampicillin and cephalexin (See Figure 3 in (3)). Scanning of its genome identified two genes coding for beta-lactamases namely *bla*_OXA-213_-like (locus tag: YA64_005895) and *bla*_ADC_ (locus tag: YA64_000855). The *bla*_oxa-213_-like gene is commonly found among members of *Acinetobacter calcoaceticus* and *Acinetobacter pittii* (Figure 2) with demonstrated resistance to ampicillin through heterologous expression in *Escherichia coli* host (23). A conserved penicillin-binding domain was identified in the *bla*_oxa-213_-like protein of LC510 which provides additional support to its role in conferring resistance to ampicillin. The resistance of LC510 to cephalexin is likely explained by the presence and expression of a *bla*_ADC_ gene encoding for AmpC beta-lactamase (24). Cloning and regulated expression of these two *bla* genes will be instructive to verify their *in-silico* predicted role in hydrolyzing beta-lactam drugs (24). The presence of *bla*_ADC_ and *bla*_OXA-213_-like genes in cave isolate LC510 that has no prior history of anthropogenic exposure supports previous work claiming that these genes contribute to the intrinsic antibiotic resistance in *A. pittii* (6, 23). The intrinsic ampicillin resistance of *A. pittii* can be suppressed with sulbactam, a *beta*-lactamase inhibitor (25), making ampicillin-sulbactam an effective antibiotic for the treatment of carbapenem-resistant *A. pittii* (7, 26).

**Figure 2.**
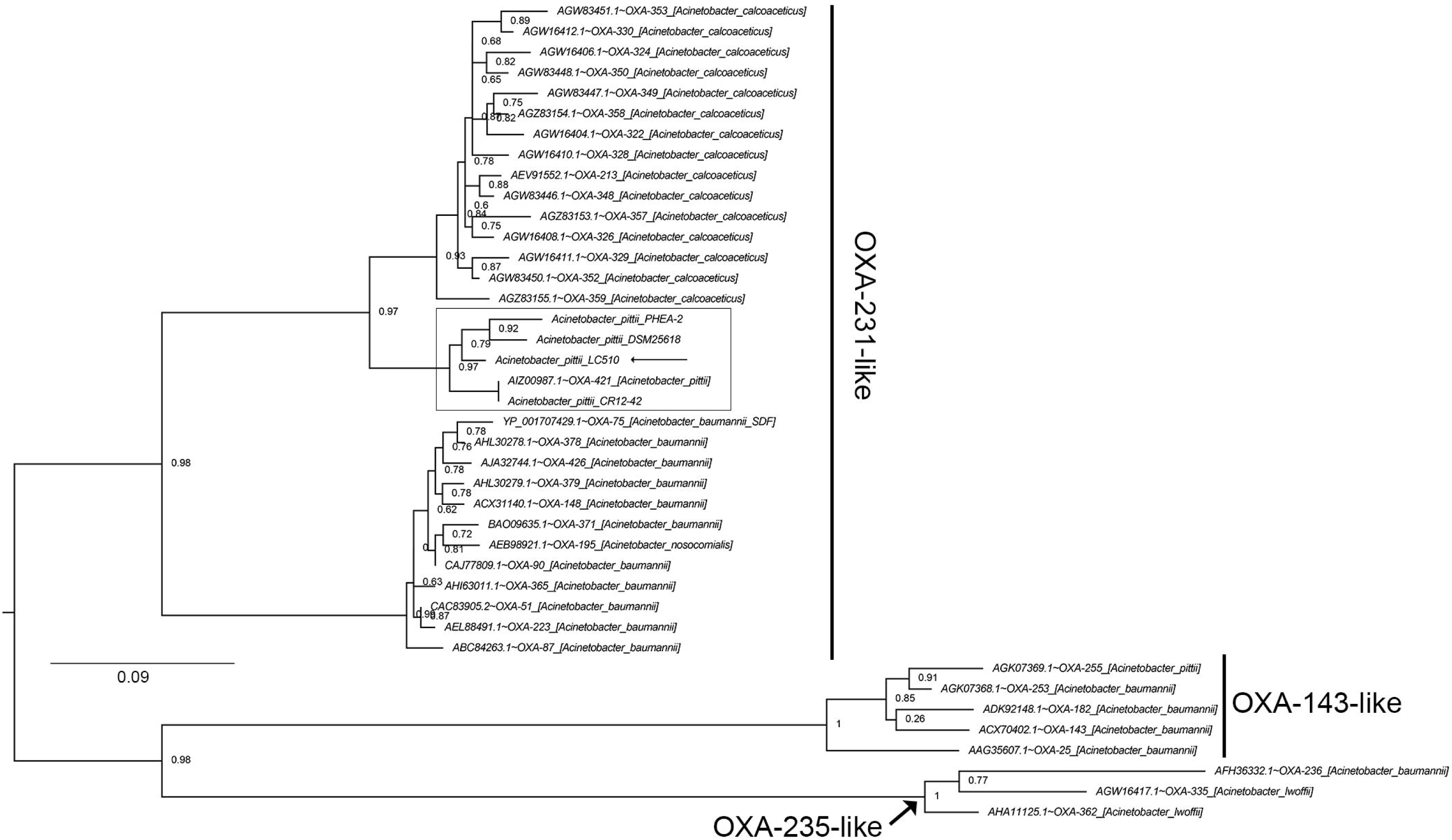
Maximum likelihood tree depicting the evolutionary relationships of *bla* proteins found in *Acinetobacter* species. Boxed clades indicate the OXA-231-like proteins from *Acinetobacter pittii* and the tree were rooted with OXA-143-like and OXA-235-like proteins as the outgroup (31). Tip labels were formatted as “GenBank Accession Number”∼ “OXA variant” _ “[strain species and ID]”. Branch lengths indicate the number of substitutions per site.

**Figure 3.**
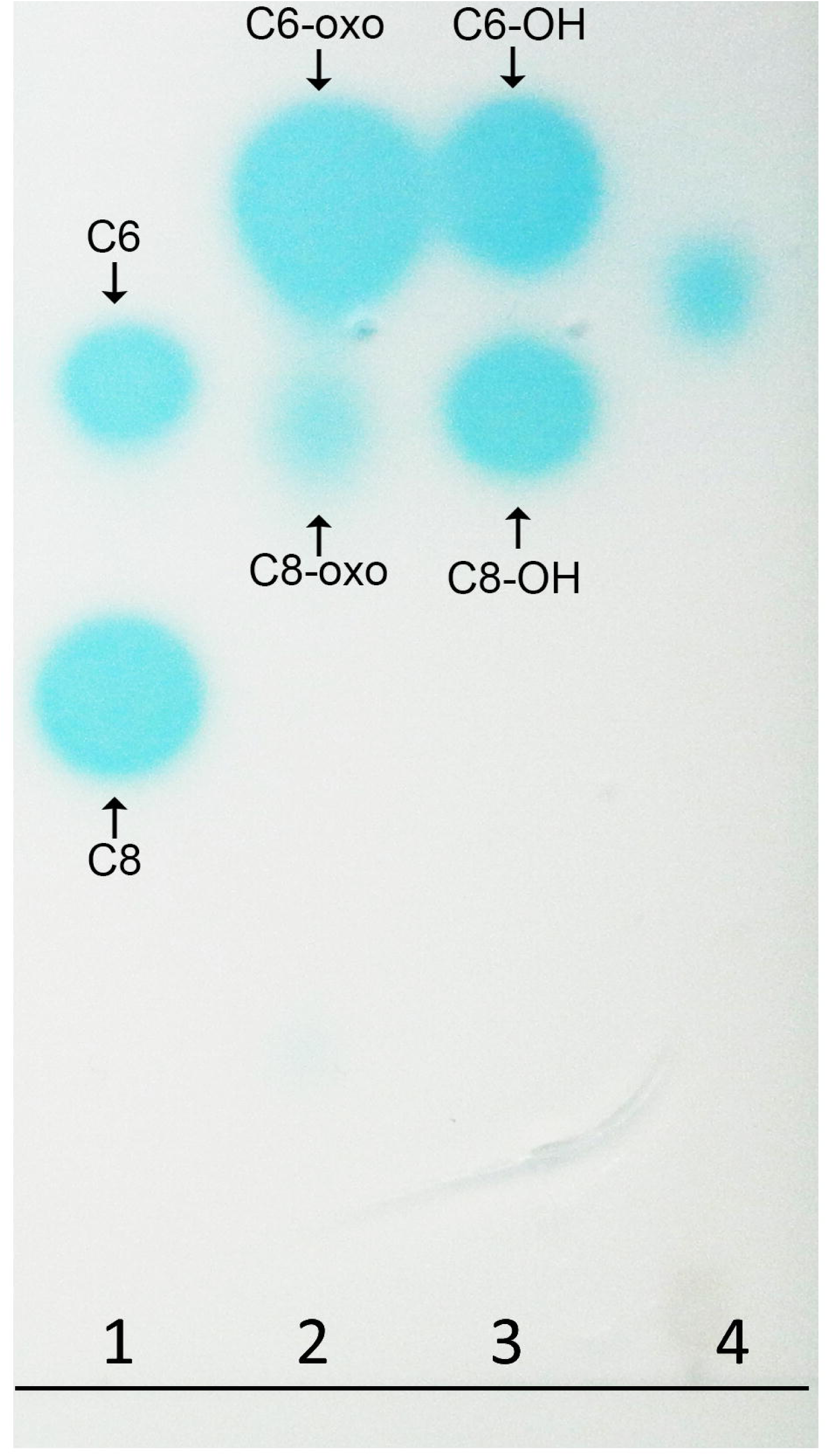
Thin Layer Chromatography of AHL signal extract of LC510 and known AHL standards separated using 70:30 Methanol:Water and overlaid with *Agrobacterium tumefaciens* biosensor in the presence of β-galactosidase substrate, X-Gal (19). Lanes 1 to 3 consist of known AHL standards while Lane 4 consists of 25 μl of 20 × ethyl acetate culture extract equivalent to 400 mL of the overnight LC510 culture. C6, N-Hexanoyl-L-homoserine lactone; C8, N-octanoyl-L-Homoserine lactone; C6-oxo, N-β-oxo-Hexanoyl-L-homoserine lactone; C8-oxo, N-β-oxo-octanoyl-L-Homoserine lactone; C6-OH, N-(3-hydroxy-hexanoyl)-L-homoserine lactone; C8, N-(3-hydroxy-octanoyl)-L-Homoserine lactone.

### Detection of quorum-sensing signal molecules and identification of an autoinducer synthase gene in *Acinetobacter pittii* LC510

Given the demonstrated ability of several *Acinetobacter* strains to produce quorum-sensing signals that are implicated in the regulation of virulence factors and cell motility, we used both *in silico* and *in vivo* approaches to assess the presence of a quorum-sensing system in strain LC510. Under the described culturing condition, LC510 strain appears to produce a medium-length AHL signal that exhibits a migration rate between C6-OH and C8-OH (Figure 3). The *luxI* and *luxR* homologs in LC510 were localized on contigs 41 and 18, respectively that exhibit strikingly high synteny to the luxI/*luxR* gene cluster in *A. pittii* PHEA-2 (Figure 4). Such a gene organization was similarly found in *Acinetobacter baumannii* M2 (27), hinting the conservation of this gene cluster and its quorum sensing (QS)-regulated genes among members of *Acinetobacter*. Transposon disruption of the *luxI* homolog (*abaI*) in strain M2 led to a substantial reduction in motility that could be rescued with the supplementation of its cognate AHL signal in the media (27). The presence of this gene cluster in LC510 may suggest that the role of quorum sensing in regulating the motility of LC510 in its cave environment. The construction of *luxI* mutant for LC510, using either transposon mutagenesis or homologous recombination (28, 29), followed by transcriptome sequencing (30) will be extremely useful not only for validating the role of QS in cell motility but also in discovering other genes and phenotypes that may be regulated by QS.

**Figure 4.**
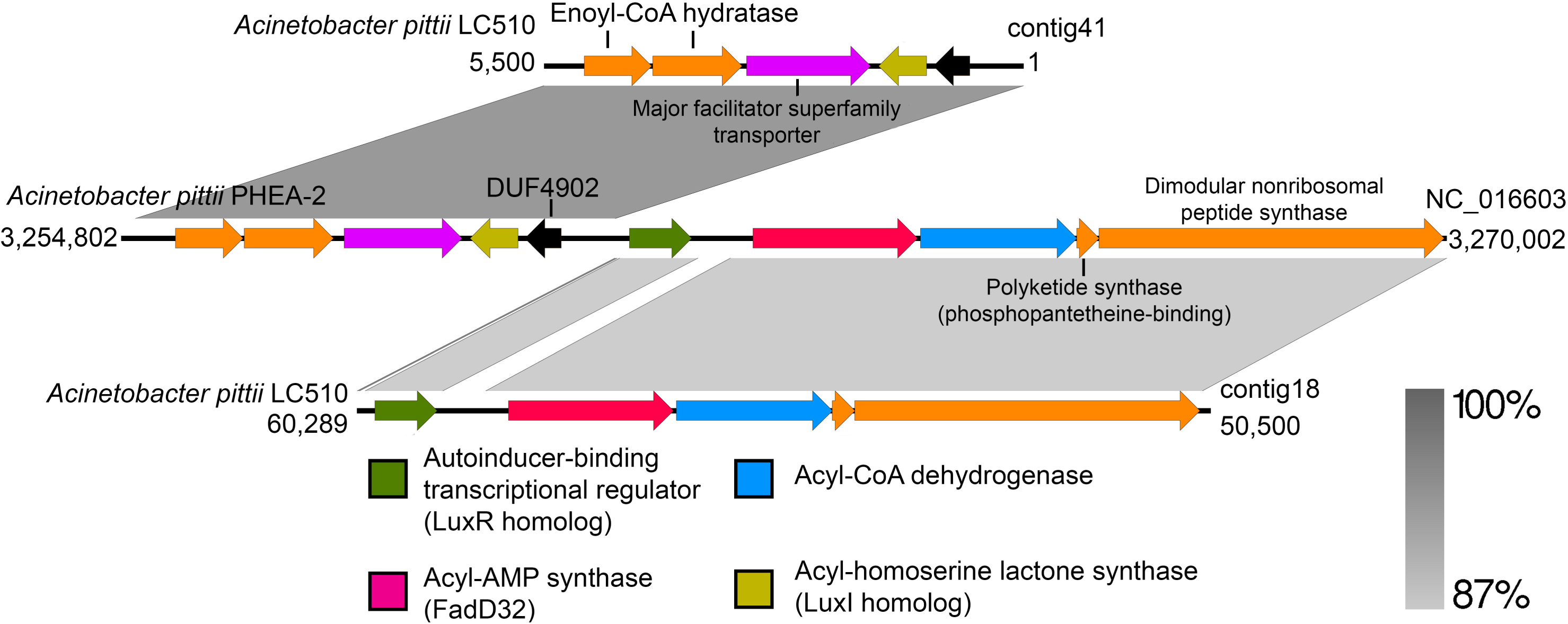
Linear genome comparison of the LC510 contigs containing the *luxR* and *luxI* homologs with closely related *Acinetobacter pittii* PHEA-2 strain. The directions of the arrows indicate transcription orientation. Note: LC510 contigs 41 and 18 contain *luxI* and *luxR* homolog, respectively.

## Conclusions

The whole-genome sequence of a Lechuguilla Cave isolate (LC510) belonging to the species *Acinetobacter pittii* was presented in this study. The identification of two *bla* genes in the annotated genome of isolate LC510 that has no prior history of anthropogenic exposure supports previous work claiming that these genes contribute to the intrinsic antibiotic resistance in members of the species *A. pittii*. In addition, LC510 still retains the ability to engage in cell-to-cell communication in an isolated cave ecosystem as evidenced by the presence of a *luxI* homolog in its genome and its ability to accumulate of N-acyl-homoserine lactones in culture medium.

## Data Availability

This Whole Genome Shotgun project has been deposited at DDBJ/ENA/GenBank under the accession LBHY00000000. The version described in this paper is <u>LBHY02000000</u>. Raw paired-end sequencing reads and sample metadata have been deposited into the NCBI public database under the BioProject ID PRJNA281683 (https://www.ncbi.nlm.nih.gov/bioproject/PRJNA281683/).

## Acknowledgments

PD, TP, AOH, and MAS acknowledge the College of Science and the Thomas H. Gosnell School of Life Sciences for support.

